# Genome Report: *De novo* genome assembly of the greater Bermuda land snail, *Poecilozonites bermudensis* (Mollusca: Gastropoda), confirms ancestral genome duplication

**DOI:** 10.1101/2025.08.21.671598

**Authors:** Stevie Winingear, Mark Outerbridge, Gerardo Garcia, Anne C. Stone, Melissa A. Wilson, P. David Polly

## Abstract

*Poecilozonites bermudensis*, the greater Bermuda land snail, is a critically endangered species and one of only two extant members in its genus. These snails are one of Bermuda’s few endemic animal clades and their rich fossil record was the basis for the punctuated equilibria model of speciation. Once thought extinct, recent conservation efforts have focused on the recovery of the species, yet no genomic information or other molecular sequences have been available to inform these initiatives. We present a high-quality, annotated genome for *P. bermudensis* generated using PacBio long read and Omni-C short read sequencing. The resulting assembly is approximately 1.36 Gb with a scaffold N50 of 44.t Mb and 31 chromosome-length scaffolds. Nearly 43 percent of the genome was identified as repeat content. This assembly will serve as a resource for the conservation and study of *P. bermudensis*, and its only close extant and also critically endangered relative, *P. circumfirmatus*. Additionally, this genome adds to the growing body of data needed for a more complete understanding of gastropod evolution and for evolutionary processes in general.

**Article summary:** This paper is aimed primarily at researchers studying genomic evolution and phylogenetics of mollusks, and secondarily researchers studying evolution and conservation of Bermuda land snails. It reports not only the first genome sequence for the Bermuda land snails, but the first sequence of any kind from the group and the first genome from the larger clade to which they belong. The paper presents a complete *de novo* genome annotation and first order analyses. The results support previous hypotheses that a genome duplication occurred at the base of land snails, and they facilitate future molecular work on Bermudian and other snails.

## Introduction

The Bermudian land snail, *Poecilozonites bermudensis*, has been a model system for the study of evolution and is now critically endangered (Gould, 1969; Outerbridge et al., 2019; Outerbridge & Sarkis, 2018). *P. bermudensis* is a surviving member of a speciose endemic radiation unique to Bermuda, a remote group of islands in the mid-Atlantic Ocean, that left an extensive fossil record that inspired the punctuated equilibrium model of evolution (Gould, 1969; Gould & Eldredge, 1977) (**Figure 1**). As many as 12 fossil species of *Poecilozonites* have been identified (Hearty & Olson, 2010), but only two remain extant, *P. bermudensis* and *P. circumfirmatus*, both of which are critically endangered (a third, *P. reinianus* became extinct sometime in the mid-20th century). *P. bermudensis* was once abundant across Bermuda, but it declined rapidly after predatory snails were introduced in the 1950s and was thought to be extinct by the early 2000s. After unexpected rediscovery in 2014, breeding colonies were established at the Chester Zoo in the UK, and some individuals have been reintroduced into protected reserves in Bermuda (Outerbridge and Sarkis, 2018; Outerbridge et al., in review). Here we present an assembled and annotated genome for *P. bermudensis and* utilize this sequence to perform comparative genomics analyses with other snail species.

**Figure 1.**
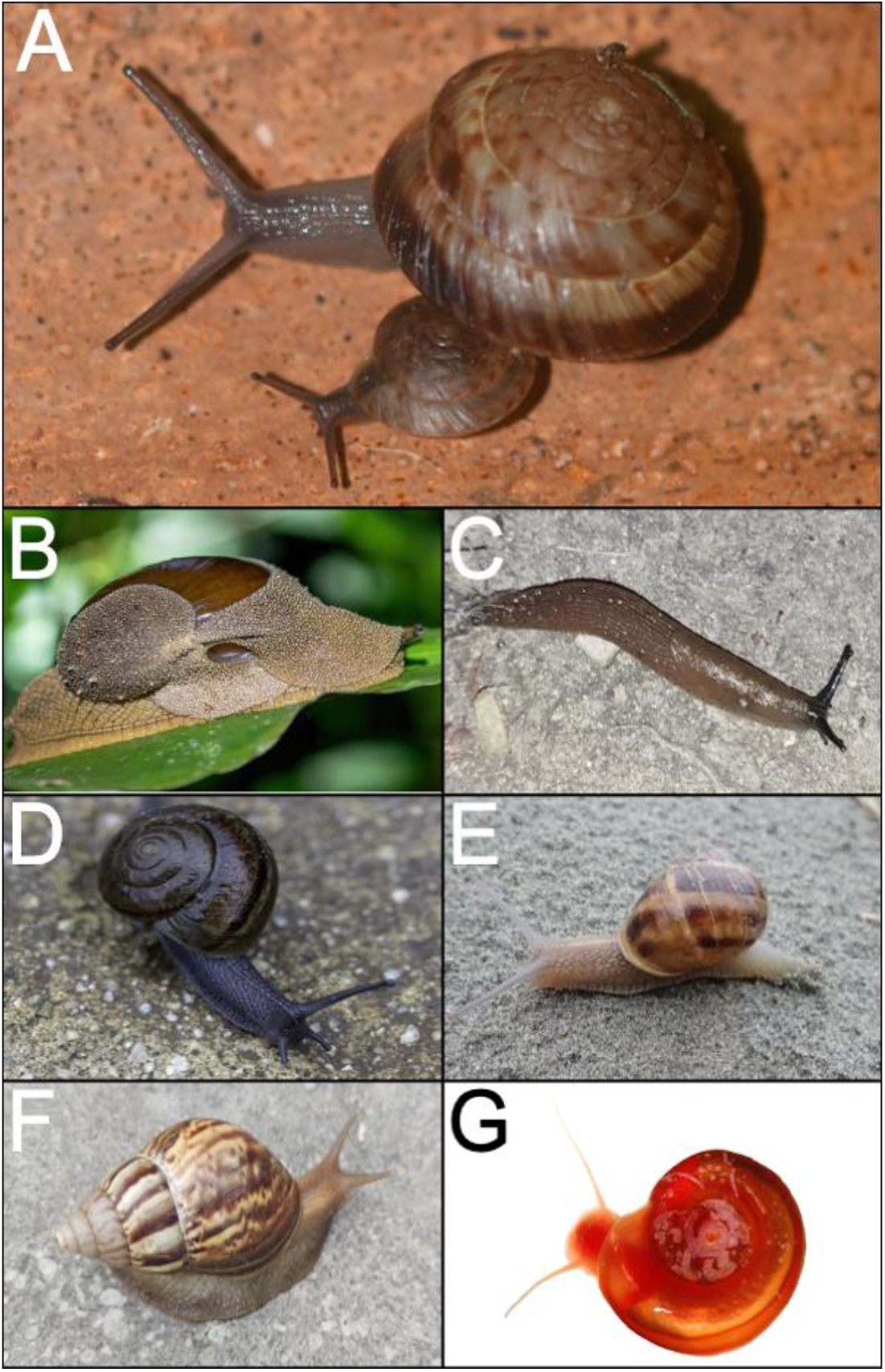
Representative snails in this study. (A) *P. bermudensis*. (B) *Megaustenia siamensis*. (C) *Arion vulgaris*. (D) *Helminthoglypta traskii*. (E) *Cornu aspersum*. (F) *Lissachatina fulica* (=*Achatina fulica*). (G) *Biomphalaria glabrata. P. bermudensis* by G. Garcia, comparative species photographs sourced from iNaturalist and Wikimedia Commons (see Supplementary Table 1).

The diversity, genetics, and evolution of gastropods are very poorly studied. After insects, gastropods rank as the second most biodiverse group, with nearly 98,000 extant species occupying oceanic, freshwater, and terrestrial habitats (McArthur & Harasewych, 2003). They are one of the few groups that made the evolutionary transition from marine to fully terrestrial environments. For example, they have developed morphological and physiological adaptations to facilitate the breathing of air and to withstand new environmental pressures like fluctuating temperatures and increased gravitational force (Vermeij & Dudley, 2000).

**Table 1.**
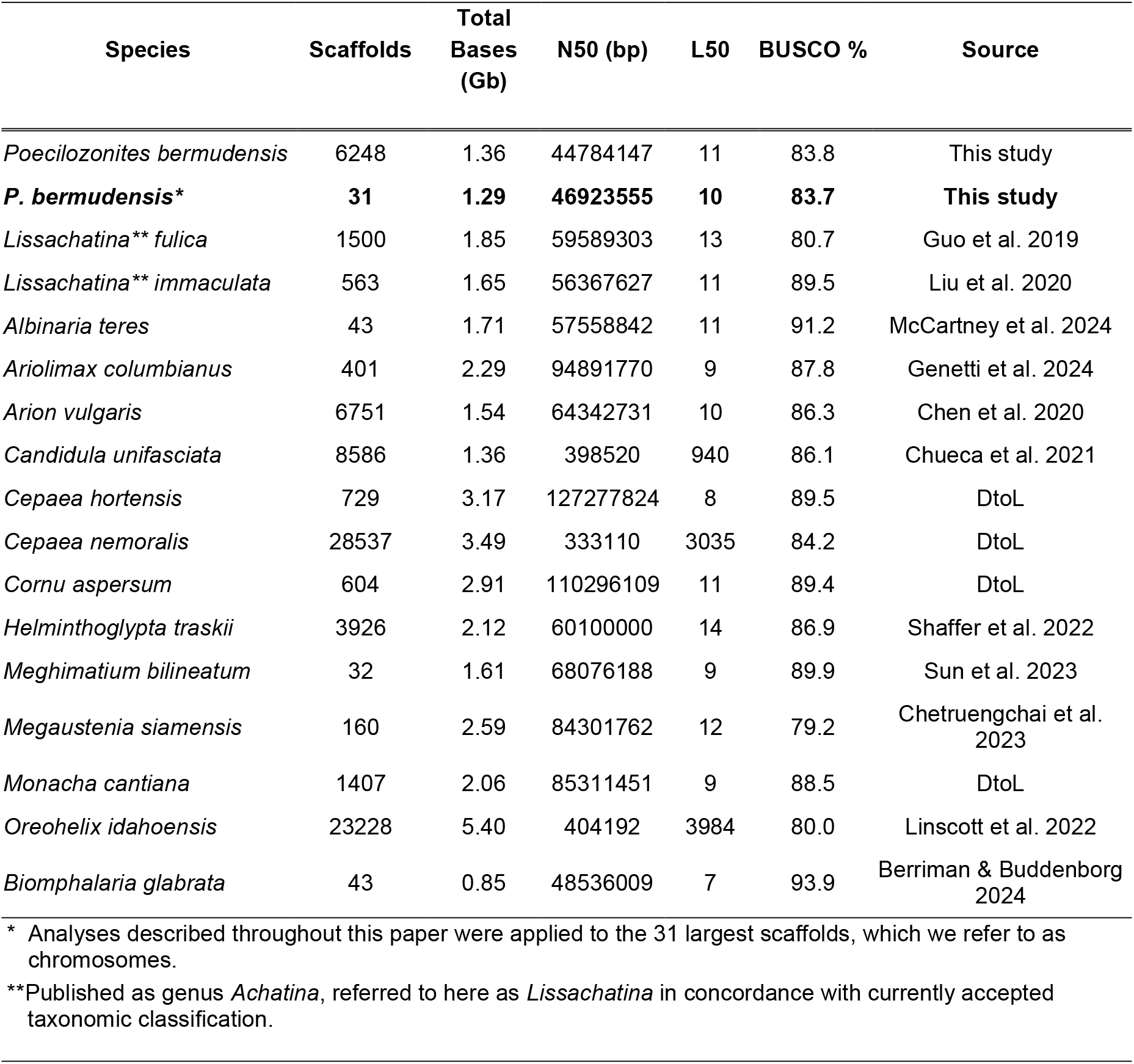
*P. bermudensis* genome assembly compared to other terrestrial snail assemblies. N50 = the length of the shortest contig such that all contigs of that length or longer cover at least 50% of the total genome length, L50 = the minimum number of scaffolds required to cover 50% of the total assembly length, BUSCO% = percentage of BUSCO genes present in the reference genome relative to the mollusca_odb10 reference dataset from OrthoDB v10.

As of 2024, the International Union for Conservation of Nature (IUCN) has classified 926 species as extinct; gastropods represent a significant portion of this with 264 extinct species listed (IUCN, 2024). Given that the IUCN’s data on invertebrates are less comprehensive than those for vertebrates, this figure likely represents a considerable underestimation of the true number of gastropod extinctions (Chiba & Cowie, 2016; Cowie & Holland, 2006; Régnier et al., 2017). Further, only a fraction of snail species have been the subject of any kind of molecular sequencing, and those that have are mostly limited to very short sequences from nuclear 18s or 28s rDNA and mitochondrial 16s rDNA genes. Fewer than 70 snail and slug species have assembled genome sequences, and those that do primarily interface directly with humans as food (such as abalone), invasive pests (apple snails), or vectors of disease (planorbid snails). Of the terrestrial subclade Stylommatophora, the land snails and slugs, fewer than 30 of its estimated 20,000 plus species have had their genomes assembled, and fewer have been carefully annotated. At present, Stylommatophora is considered a monophyletic clade divided into two groups: “achatinoid” and “non-achatinoid”, based on mitochondrial (16s, cytochrome c oxidase I) and nuclear (18S, 28S) markers (Ayyagari & Sreerama, 2020; Guzmán et al., 2021; Wade et al., 2001, 2006). Understanding these taxonomic relationships is essential for studying land snail evolution, but the limited availability of high-quality genome assemblies continues to prohibit deeper phylogenetic and other evolutionary analyses.

The genome assembly of *P. bermudensis* opens new possibilities for understanding processes of speciation in this unique, isolated endemic radiation of snails and provides critical resources for conservation and breeding efforts to prevent its extinction. Adding to the pool of high-quality whole genome sequences in terrestrial gastropods, we help to advance our understanding of the evolution of terrestriality, phylogenetics, and conservation efforts across this extensive clade.

## Materials and Methods

### Sample collection, DNA preparation and sequencing

Three *P. bermudensis* juveniles from the breeding colony at Chester Zoo (Upton-by-Chester, Chester, CH2 1EU United Kingdome; Registered Charity No. 306077) were flash frozen in liquid nitrogen and provided to Dovetail Genomics (now Cantata Bio) (Scotts Valley, CA) where DNA extraction, library preparation, sequencing, assembly, and annotation were performed. Approximately 200-500 mg of foot tissue from one individual was used for this assembly. Tissue collection was performed in accordance with the UK Animals (Scientific Procedures) Act 1986 under the Chester Zoo ethics number 202101-01 (approved 04 January 2021).

High molecular weight DNA was extracted from the tissue and then prepared for sequencing by Dovetail staff. First, continuous long-read libraries were created and sequenced using PacBio technology. A short read library of chromatin conformation was produced using Dovetail’s proprietary Omni-C (or “Chicago”) library preparation method (Putnam et al., 2016). In brief, the Omni-C libraries are chromatin fixed and crosslinked to preserve information about long (and short) range associations within a genome, and they improve the scaffolding of de novo assembled genomes created using high-throughput sequences. Short read libraries were sequenced to approximately 30X coverage using the Illumina HiSeqX platform.

### Genome assembly and annotation

Long read PacBio sequences were assembled using wtdbg2 v2.5 (Ruan & Li, 2020). Short read Omni-C sequences, along with the initial PacBio assembly were used as input for the HiRise pipeline (described by Putnam et al., 2016) to generate longer, more continuous scaffolds.

Initial genome annotation was completed using the Dovetail Genomics genome annotation workflow as detailed in Jarvis et al. (2017). Briefly, repeat families found in the genome assemblies of *P. bermudensis* were identified de novo and classified using the software package RepeatModeler (v 2.0.1, Flynn et al., 2020) and masked using RepeatMasker (v 4.1.0; Smit, Hubley & Green, 2013-2015). Publicly available coding sequences from *Lissachatina immaculata* and *Pomacea canaliculata* (Guo et al., 2019; Liu et al., 2018) were used to train the initial ab initio model for *P. bermudensis* using both the AUGUSTUS software (v 2.5.5; Stanke et al., 2008) and SNAP (v 2006-07-28; Korf, 2004). RNAseq reads were mapped onto the genome using the STAR aligner software (v 2.7; Dobin et al., 2013) and MAKER (Cantarel et al., 2008), SNAP and AUGUSTUS (with intron-exon boundary hints provided from RNA-seq) were then used to predict genes in the repeat-masked reference genome. Only genes that were predicted by both SNAP and AUGUSTUS softwares were retained in the final gene set.

To assess the completeness of the *P. bermudensis* assembly, BUSCO analysis was performed using BUSCO v 5.7.1, referencing the most recent mollusk specific BUSCO dataset - in this case mollusca_odb10 (Manni et al., 2021a,b).

### Phylogenetics Analysis

Phylogenetic analyses were conducted to assess relationships within Stylommatophora, with a particular focus on clarifying the placement of *P. bermudensis* relative to other members of the order. All stylommatophoran snail and slug species with fully assembled genomes available at the time of writing were included. *Biomphalaria glabrata*, an aquatic pulmonate, was included as an outgroup.

Using the custom pipeline available at https://github.com/jamiemcg/BUSCO_phylogenomics (McGowan, 2023), we identified 936 single copy orthologs shared by all analyzed species. These genes were aligned individually using MUSCLE (Edgar, 2004), and the resulting alignments were trimmed using trimAL. Within this pipeline, we employed TrimAL’s default setting, allowing for selection of an optimal trimming method based on average identity scores and total number of sequences in the alignment (Capella-Gutierrez et al., 2009). Maximum likelihood trees for each gene were then inferred using IQ-TREE (Minh et al., 2020) and ASTRAL-IV (Zhang & Mirarab, 2022) was implemented to estimate the species tree by identifying the complete phylogeny that contained the most shared topology with the given set of gene trees. Branch support for the final phylogeny was calculated in ASTRAL-IV as local posterior probabilities (Sayyari and Mirarab, 2016).

### Synteny Analysis

Evidence for whole genome duplication events in the evolutionary history of *P. bermudensis* was assessed through synteny analysis. Using 936 single copy orthologs identified by BUSCO, we compared the chromosomal arrangement of *P. bermudensis* to other terrestrial snails and slugs, as well as to *Biomphalaria glabrata*, a marine species outside of the stylommatophoran clade. BUSCO’s accompanying support protocol was used to map the positions of each identified ortholog and infer syntenic relationships between genomes (Manni et al., 2021b).

## Results and Discussion

### Genome assembly and annotation

Using PacBio continuous long reads (555 gigabase-pairs, 278X coverage) and Illumina short reads (58.9 million read pairs, 30X coverage), we assembled the genome of *P. bermudensis*. The resulting draft genome assembly is 1.36 Gb in size, comprising 6248 scaffolds, with a scaffold N50 of 44.7 Mb. Ambiguous bases, Ns, account for only 0.10% of the assembly, demonstrating high continuity and quality. Repeat analysis using RepeatModeler indicated approximately 42.9% of the genome consists of repetitive elements, primarily Class I retroelements (13.5%, the majority of which are LINEs), and Class II DNA transposons (3.77%). Simple repeats (3.47%) and low-complexity regions (0.37%) were also identified, and the remaining 21.7% of repeats remain unclassified.

Genome annotation predicted 39,119 genes, comprising 44.3 Mb of coding sequence, with an average gene length of 1,132 bp. An evaluation of completeness using BUSCO analysis (reference database mollusca_odb10) reveals the complete presence of 83.8 percent of expected genes. BUSCO analysis is useful for evaluating genome quality as it checks for essential genes expected to be present in a complete genome based on a particular database.

After scaffolding, we identified 31 scaffolds likely representing chromosome-length sequences (**Figure 2**). When BUSCO analysis was run specifically on these 31 chromosome-length scaffolds, the completeness percentage was 83.7. This close match increases our confidence that the identified scaffolds represent chromosomes and contain the majority of the genome assembly. These BUSCO results are comparable in quality to other recently produced stylommatophoran genomes (**Table 1**).

**Figure 2.**
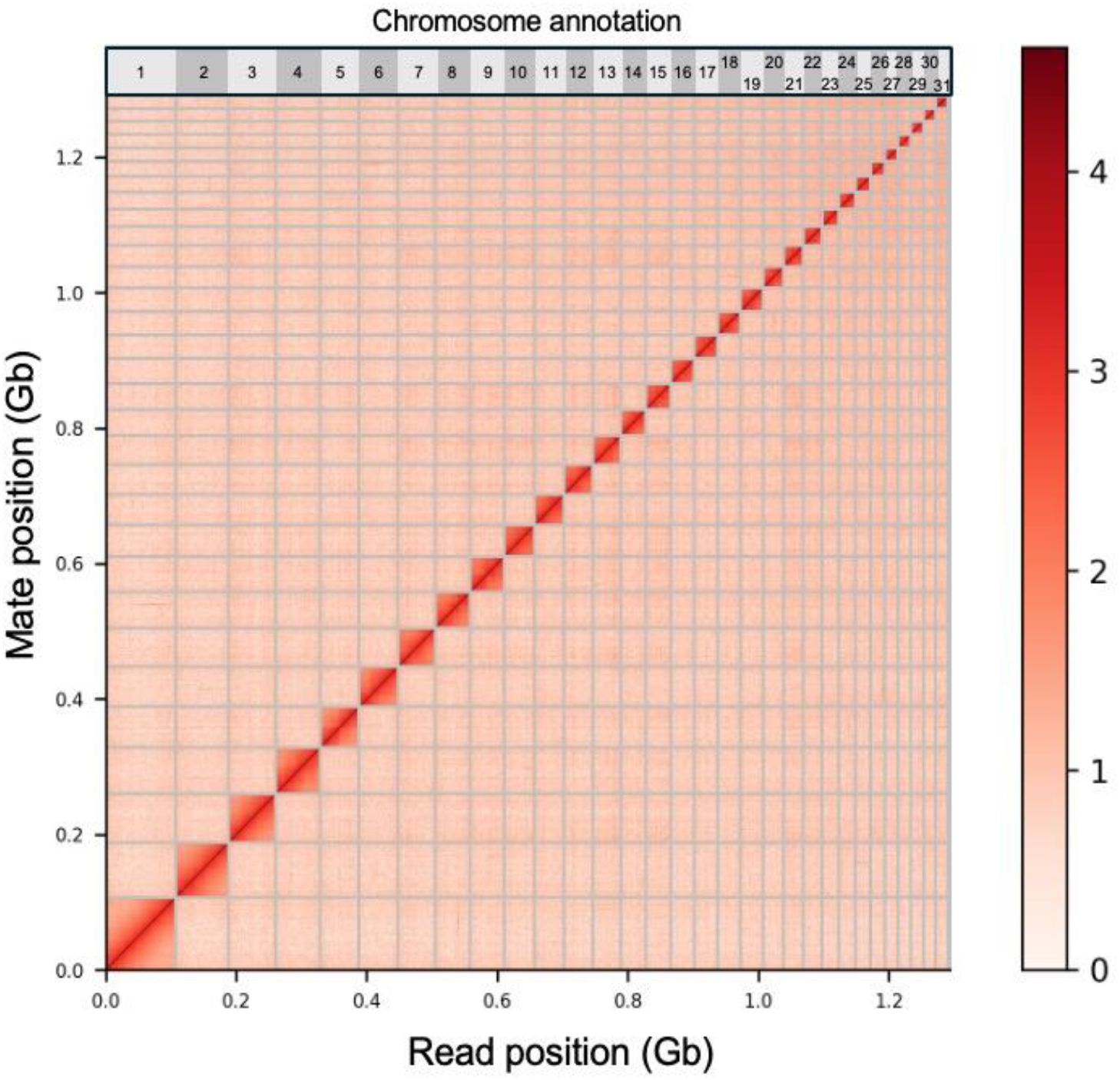
HiC Contact map. HiC chromatin conformation capture map across the 31 largest scaffolds, which we annotate from largest to smallest as putative chromosomes in *P. bermudensis*.

### Phylogenetic analysis

We used the resulting phylogeny to assess the placement of *P. bermudensis* within those 15 stylommatophoran species with assembled genomes (**Figure 3**). The two achatinoid species clustered together into a distinct clade. Among the non-achatinoid species, *P. bermudensis* was most closely related to *Megaustenia siamensis*, a snail species that is endemic in southeast Asia, albeit with a divergence time that could still be quite deep because so few snail groups have yet been sequenced. These two cluster with the three available slug genomes, and then with *Albinaria teres*, a mediterranean snail species found primarily in Greece, and *Oreohelix idahoensis*, a snail native to the western United States. These species collectively form a clade that is sister to one containing primarily western European snails.

**Figure 3.**
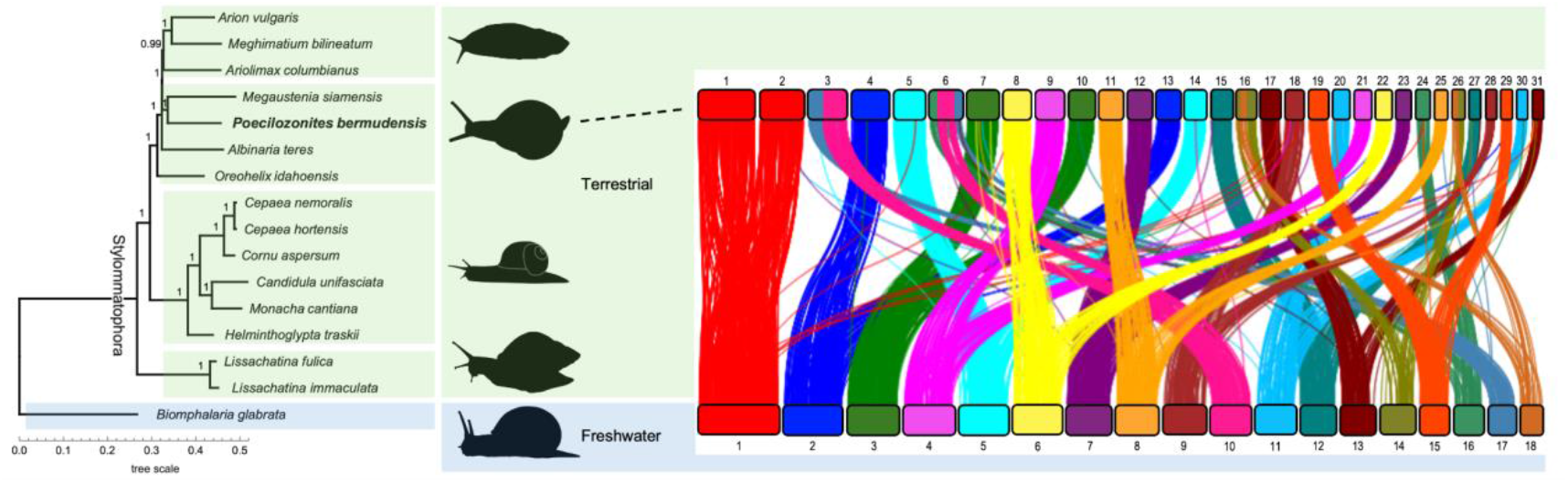
Phylogenetic relationships of *Poecilozonites bermudensis* and synteny with *Biomphalaria glabrata*. Species tree (left) is a consensus of 936 approximately-maximum-likelihood gene trees based on all shared single copy BUSCO genes. Branch support values are local posterior probabilities (all branches ≥0.99). Branch lengths correspond to substitutions per site. The synteny plot (right) illustrates the chromosomal arrangement of BUSCO genes in *P. bermudensis* and *B. glabrata*, where lines represent orthologous loci. Silhouettes from PhyloPic (https://www.phylopic.org; Keesey, 2025).

The topology recovered here is consistent rDNA trees for the five species common to previous studies (Wade et al. 2001) and confirms that *Poecilizonites* is part of the Limacoidea clade. Our results are also consistent with older phylogenetic suppositions based on comparative anatomy that suggested *Poecilozonites* is closely related to gastrodontid snails (Pilsbry, 1924; Hausdorf, 1998). Bermuda has never been connected to any other landmass (Coates et al., 2013), suggesting that the ancestors of *Poecilozonites* likely reached the island in small numbers from North America. To improve on existing inferences about the timing and route of colonization, future research should focus on sequencing a broader and more geographically diverse array of stylommatophoran genomes. Special attention should be given to species from regions closest to Bermuda, particularly those along the southeast coast of the United States, which may shed light on potential source populations.

### Whole genome duplication

Whole genome duplications (WGDs) have long been hypothesized to play a crucial role in the evolutionary history of molluscs. Evidence of natural polyploidy among molluscs Goldman et al., 1983; Thiriot-Quiévreux, 2003), and the feasibility of inducing polyploidy experimentally (Cadoret 1992) suggests that genome duplication has significantly influenced molluscan adaptation. Such paleoploidy events have been proposed in the cephalopods (Hallinan and Lindberg, 2011; Yoshida et al., 2011) and stylommatophorans (Hallinan and Lingberg, 2011).

More recent molecular investigations have modified Hallinan and Lingberg’s hypothesis, instead suggesting that chromosomal rearrangements contributed more significantly to deeper molluscan evolution than genome duplication (Albertin et al, 2015 and 2022, Jiang et al 2022; Belcaid 2019). However, a WGD likely did occur somewhere in the stylommatophoran lineage. This is supported by whole genome synteny analyses of the stylommatophoran species *A. vulgaris* (Chen et al., 2022) and giant African land snails (Guo et al., 2019; Liu, 2020). Further evidence to support the occurrence of this WGD comes from our analysis of orthologous gene synteny among land snails.

Our analysis shows that when compared to a non-stylommatophoran outgroup, *B. glabrata*, genes from a single chromosome appear to share homology across two chromosomes in *P. bermudensis* (**Figure 3**). Notably, our analysis also suggests synteny of gene content is extremely well conserved; genes from one *B. glabrata* chromosome map across two chromosomes from *P. bermudensis* with little gene homology outside of this pairwise relationship.

The precise timing and location of the stylommatophoran WGD as hypothesized by Hallinan and Lindberg remains undetermined (Hallinan & Lindberg, 2011). They suggested that this event might have occurred early in the lineage, either at the base of the entire branch—potentially facilitating the transition of mollusks to terrestrial environments—or slightly later at the base of what they define as the Sigmurethra-Orthurethra branch (taxon names that have since been abandoned) following divergence from the Succineidae. Historically, succineids were considered a sister group to Stylommatophora; however, these more recent analyses have placed succineids within the non-achatinoid lineage of Stylommatophora.

Currently, no complete genomes have been assembled for any succineid species. Available karyotype data indicate that these species generally have fewer chromosomes—commonly less than 10, although variability exists with some having 20-25 chromosomes (Barker, 2001; Kiauta and Butot, 1968). The evidence of WGD in both giant African land snails (achatinoid) and *P. bermudensis* (a non-achatinoid species) suggests that the WGD might have occurred at the base of Stylommatophora. Nevertheless, the notably smaller number of chromosomes in succineids compared to other groups raises questions about this hypothesis. Until genomes from some succineid species can be included in syntenic analyses, the exact placement of this WGD remains speculative.

## Conclusions

This study presents the draft assembly of *Poecilozonites bermudensis*, achieving an assembly quality level commensurate with other published land snail genomes. The availability of this genome represents an essential first step in advancing research on Bermuda’s land snails, laying the groundwork for future conservation efforts and incorporating the species into the broader body of literature concerning the evolutionary history of gastropods.

## Data availability

Raw PacBio and HiC sequence data are through the National Center for Biotechnology’s (NCBI) database under Bioproject PRJNA1250146. The complete assembled *P. bermudensis* genome was deposited under the accession JBNFNQ000000000 - the assembly described in this paper is version JBNFNQ010000000. Genome annotation files have been deposited in Figshare along with all code used in these analyses (DOI 10.6084/m9.figshare.28791785).

## Acknowledgements

The tissue was imported to the US with US Fish & Wildlife Service Permit 70881B to Dovetail Genomics, LLC. Tissue was harvested in accordance with Chester Zoo’s Animal Subject Ethics guidelines.

Thanks to Rebecca Mogey at Chester Zoo for extracting and shipping the tissue; Jasmine Haimowitz and Joanna Gallagher at Dovetail Genomics for coordinating the shipping; Jordan Zhang for work on the annotation; Thomas Swale and Mark Daly at Dovetail for assistance with processing and funding; Gieslla Caccone and Kirsten Dion at Yale University for suggestions about DNA extraction. We also thank the Darwin Tree of Life Project for access to genomic resources used for comparative analysis. The authors acknowledge Research Computing at Arizona State University for providing high performance computing resources that have contributed to the research results reported within this paper.

## Funding

The genome assembly was generated as part of a Tree of Life Grant from Dovetail Genomics. This publication was supported by the National Institute of General Medical Sciences of the National Institutes of Health under Award Number R35GM124827 to MAW. This work was supported, in part, by the Intramural Research Program of the National Human Genome Research Institute, National Institutes of Health.

This research was supported in part by the Intramural Research Program of the National Institutes of Health (NIH). The contributions of the NIH author(s) are considered Works of the United States Government. The findings and conclusions presented in this paper are those of the author(s) and do not necessarily reflect the views of the NIH or the U.S. Department of Health and Human Services.

